# Chemosynthesis enables microbial communities to flourish in a marine cave ecosystem

**DOI:** 10.1101/2025.10.22.684061

**Authors:** Francesco Ricci, Tess Hutchinson, Pok Man Leung, Thanh Nguyen-Dinh, Jialing Zeng, Thanavit Jirapanjawat, Vera Eate, Wei Wen Wong, Perran L.M. Cook, Chris Greening

## Abstract

Chemosynthesis, an ancient metabolism that uses chemical compounds for energy and biomass generation, occurs across the ocean. Although chemosynthesis typically plays a subsidiary role to photosynthesis in the euphotic ocean, it is unclear whether it plays a more important role in aphotic habitats within this zone. Here, we compared the composition, function, and activity of sedimentary microorganisms within a marine cave at mesophotic depth, across a transect from the entrance to the interior. Microbes thrived throughout this ecosystem, with interior communities having higher diversity than those at the entrance. Analysis of 132 species-level bacterial, archaeal, and eukaryotic metagenome-assembled genomes revealed niche partitioning of habitat generalists distributed along the cave, alongside specialists enriched across its entrance and interior environments. Photosynthetic microbes and photosystem genes declined in the inner cave, concomitant with enrichment of chemosynthetic lineages capable of using inorganic compounds such as ammonium, sulfide, carbon monoxide, and hydrogen. Biogeochemical assays confirmed that the cave communities consume these compounds and fix carbon dioxide through chemosynthesis, with inner communities mediating higher cellular rates. Together, these findings suggest that the persistent darkness and low hydrodynamic disruption in marine cave sediments create conditions for metabolically diverse communities to thrive, sustained by recycling of inorganic compounds, as well as endogenous and lateral organic matter inputs. Thus, chemosynthesis can sustain rich microbial ecosystems even within the traditionally photosynthetically dominated euphotic zone.

## Introduction

Chemosynthesis, the microbial metabolic process that transfers carbon to the biosphere using chemical energy, provides a foundation for marine life^1^. Once thought to be limited to hydrothermal vents and other geologically active environments^2–6^, chemosynthesis is now recognised as far more pervasive throughout the ocean^1^. For example, activity-based experiments have shown that surface ocean microbial communities oxidise ammonia (NH_4_^+^), carbon monoxide (CO), and molecular hydrogen (H_2_) for both survival and growth, while co-existing with conventional photoautotrophs and organoheterotrophs^7,8^. Chemosynthesis is also dominant in certain near-surface waters where either photosynthesis is excluded, for example in waters shielded by Antarctic ice shelves^9^, or where chemical inputs are high, for example in mesophotic cold seeps where benthic fauna form mutualistic relationships with chemosynthetic microbial communities^10^, underscoring the ecological importance of this process even at relatively shallow depths. Across the global ocean, chemical energy sources not only arise from geological seepage but also from a suite of biotic and abiotic processes that generate compounds capable of fuelling chemosynthesis; for example, processes such as sulfate reduction, nitrogen fixation, and organic carbon degradation generate significant amounts of sulfide (S^2-^), H_2_, NH ^+^, and CO that can be used as energy sources for chemosynthetic aerobes and anaerobes^7,11–16^. Beyond shaping local trophic chains, chemosynthesis influences global elemental cycles, forming a foundation for marine biogeochemistry and ecology^1^. These findings, among others, have transformed our understanding of the breadth of chemosynthetic microorganisms and suggest other chemosynthetically-supported ecosystems remain to be characterised.

The vastness of the ocean and the inaccessibility of many of its environments continue to challenge our understanding of the full breadth of ecological niches and their implications for life on Earth. Among the most enigmatic are marine caves, benthic habitats found along coastal margins worldwide that form unique interfaces between the subsurface and the coastal ocean. Despite their global distribution17–20, the microbiology of marine caves remains largely unexplored, often obscured by logistical challenges in accessing them. At Cape Palinuro, Italy, fluids enriched in hydrogen sulfide (H2S), methane (CH4), thiosulfate (S2O32−), and carbon dioxide (CO2) emitted from an active spring in the Grotta Azzura sustain Beggiatoa-like microbial mats21, reminiscent of those found at abyssal vents22,23, highlighting surprising parallels between shallow and deep-sea chemosynthetic ecosystems. Other marine caves lack active vents, but experience severely restricted water exchange, resulting in periodical anoxic conditions accompanied by substantial accumulation of H2S throughout the cave water column and sediment24. Anoxygenic phototrophs, such as purple sulfur bacteria, can benefit from these conditions and proliferate in dense populations25. In these perennially light-limited systems, sulfide availability rather than light can become the primary constraint on biomass production24. These foundational studies position marine caves as environments with diverse but distinctive microbiology and biogeochemistry. Yet, no study to date has systematically investigated the composition or function of the microorganisms within marine caves. A mechanistic understanding of the ecological strategies employed by cave-dwelling microbes, the diversity of energy sources employed, and the carbon fixation pathways sustaining primary production remains limited. Equally unresolved are the bottom-up metabolic interactions that structure microbial communities and underpin biogeochemical cycling in these complex systems.

Here we address these knowledge gaps by investigating a marine cave at a depth of approximately 30 metres in Port Phillip Bay, Victoria, Australia. This site served as a natural system to explore the spatial transition of microbial metabolisms from the light-exposed, hydrodynamically active sediment at the cave entrance to the dark, stable cave interior. We hypothesised that diminishing light toward the inner cave limits phototroph abundance, whereas the reduced water flow of the interior cave facilitates the formation of stratified microbial communities, where chemosynthetic processes are fostered by interactions between anaerobic and aerobic metabolisms. Bridging gene- and genome-resolved metagenomics with *ex situ* activity incubations and isotope geochemistry, we shed light on the ecological strategies of cave-dwelling microbes and show that genes mediating the oxidation of inorganic compounds are highly abundant in the innermost cave sediments and sustain a diversity of carbon fixation pathways, which ultimately enhance the primary productivity of the cave ecosystem.

## Methods

### Sample collection, processing and physicochemical parameters

The study site was a granitic marine cave located at a maximum depth of ∼32□m in Port Phillip Bay, Melbourne (38.293733° S, 144.625867° E; **Fig. 1a, b**). Sampling during SCUBA diving at this depth for approximately 30 minutes requires strict decompression procedures to ensure divers’ safety, making fieldwork logistically challenging. Additionally, as the cave lies near an active shipping channel, access is restricted to brief windows of less than 1 hour during slack water at the end of an ebb tide and in the absence of vessel traffic. These constraints inherently limited the achievable sample size. We collected triplicate sediment samples over two expeditions in November 2023 and October 2024 along a transect extending from the cave entrance (0 m) through the midsection (6.5 m) to the innermost accessible environment (13 m), and two seawater samples, one from each extremity of the cave (**Fig. 1c**). We use the collective term interior to refer to the cave midsection and innermost accessible environment. After collection, the samples were transported to Monash University. Samples for molecular, physicochemical, and stable isotope analyses were stored in a -20 °C freezer until processing, whereas samples for respiration, trace gas consumption, nitrification, and radioisotope analyses were processed within a day. Sediment physicochemical parameters (pH, electrical conductivity, total carbon, total organic carbon, nitrate-N, ammonium-N, and sulfur) were measured at the Environmental Analysis Laboratory (EAL), Southern Cross University, NSW, Australia.

**Figure 1.**
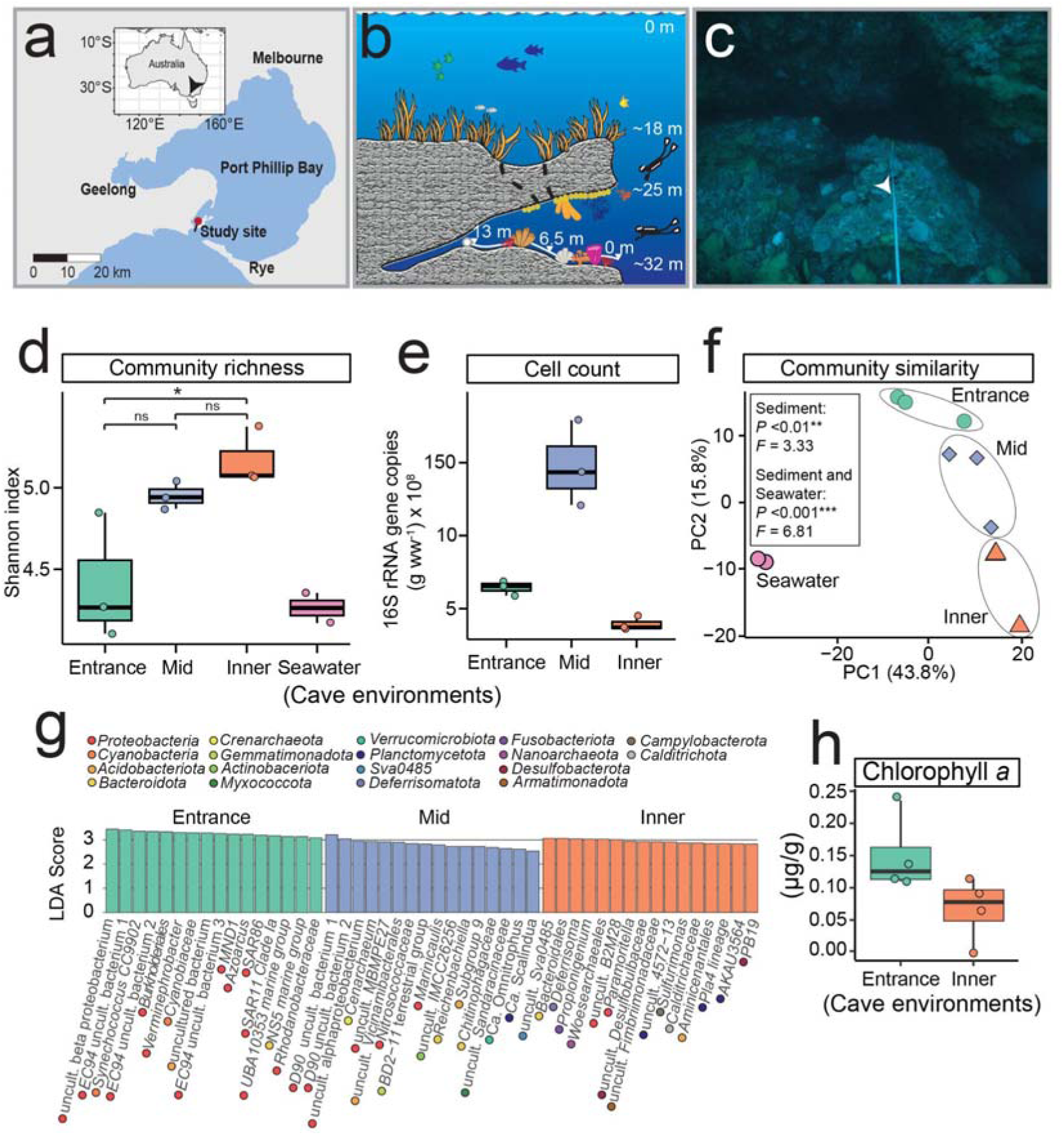
(**a**) Map of Port Phillip Bay indicating the location of the study site. (**b**) Schematic illustration of the surveyed marine cave, showing representative depths, transect across the entrance (0 m), mid (6.5 m), and inner (13 m) cave environments, and typical benthic assemblages, including cnidarians, sponges, and algae within and surrounding the cave. (**c**) Close-up of the measuring transect highlighting the cave floor substrate. (**d**) Community richness (Shannon index) of archaeal and bacterial communities in sediment and seawater, based on 16S rRNA gene data. (**e**) 16S rRNA gene copy number per gram wet weight (g ww) of sediment across cave environments. (**f**) Principal component analysis (PCA) of beta diversity showing compositional differences in sediment and seawater microbial communities. (**g**) LEfSe analysis identifying archaeal and bacterial genera enriched in entrance, midsection, and inner cave environments, with linear discriminant analysis (LDA) scores indicating discriminative power. Only top 16 enriched genera per environment are shown, for a full list refer to Supp. Data 1. (**h**) Chlorophyll *a* concentration in sediments at the entrance and inner cave entrance, indicating abundance of oxygenic phototrophs.

### Community DNA extraction and sequencing

Genomic DNA was isolated from each sediment (*n* = 9) and sea water (*n* = 2) sample using the FastDNA Spin Kit for Soil (MP Biomedical, California, USA) according to the manufacturer’s protocol, and following guidelines for preventing contamination^26^. For sediment, 0.5 g of material was used per extraction; for seawater, DNA was isolated from biomass collected on 0.22□µm filters after filtering 1.5□L of seawater. A no-template extraction control was processed in parallel. DNA libraries were sent to the Genomics Node of the Monash Genomics & Bioinformatics Platform (Melbourne, Victoria) for library preparation and sequencing on a single lane of an Illumina NovaSeqX Plus using XLEAP-SBS chemistry (2 x 150 bp).

### Reads quality control, assembly, and binning

Shotgun metagenomic sequencing of libraries generated from three biological replicates per cave environment yielded an average of 58.8□ ±□ 5.8 million reads for the cave entrance, 54.3□ ±□ 5.1 million reads for the midsection, and 53.9□ ±□ 2.5 million reads for the innermost samplable region. Seawater samples generated 8.4 and 52.7 million reads, while the extraction control yielded a total of 4,308 reads. Raw sequencing data were processed through the Metaphor pipeline27. Quality control was carried out using fastp28, which included the removal of adapters and low-quality sequences, as well as filtering by read length (≥50 bp) and average quality score (≥30), with automatic adapter detection enabled. To enhance genome recovery, particularly of low-abundance taxa, and minimise redundancy, reads from all libraries were co-assembled using MEGAHIT v1.2.929 using default parameters. Assemblies were filtered to retain contigs ≥1,000 bp. Genome binning was independently performed with four algorithms: Vamb v4.1.330, MetaBAT v2.12.131, CONCOCT v1.1.032, and SemiBin v2.2.033.

### Bacterial and archaeal bin processing

Bins were consolidated using DAS Tool v1.1.634 to generate a refined bin set. These bins were subsequently dereplicated with dRep v3.4.235 at a 95% average nucleotide identity (ANI) threshold corresponding to species level, incorporating CheckM236 metrics. Completeness and contamination were assessed with CheckM236. In total, we recovered 130 bacterial and archaeal metagenome-assembled genomes (MAGs) meeting the MIMAG criteria37 for medium-(≥50% completeness; <10% contamination) and high-quality (≥90% completeness; <5% contamination). MAGs were taxonomically classified using GTDB-Tk v2.3.238 against the Genome Taxonomy Database (GTDB) release R22039. The coverage of each MAG across samples was calculated using CoverM v0.7.040 in genome mode with method count. We defined seawater-exclusive MAGs as those with less than 1,000 reads present in the sediment and/or with a 100-fold increase in the seawater relative to the sediment. A total of five MAGs met these criteria. The maximum-likelihood phylogenetic tree of Archaea and Bacteria MAGs was built through PhyloPhlAn v3.1.141 using the LG substitution matrix with diversity high, then plotted using iTOL v742 and edited in Illustrator v24.0.2.

### Metabolic annotation of metagenomic short reads and contigs

Adapter and barcode sequences were removed from raw reads, followed by the elimination of PhiX contaminants and low-quality sequences (minimum Phred score of 20) using the BBDuk utility within BBTools v36.92 (https://sourceforge.net/projects/bbmap/). Forward reads that passed quality filtering and were ≥130 bp in length were screened for functional genes using DIAMOND blastx43 against a curated and regularly updated database44 comprising 62 metabolic marker genes. These genes represent key pathways involved in energy conservation, carbon fixation, phototrophy, and the cycling of hydrogen, carbon monoxide, methane, sulfur, nitrogen, and iron. Search parameters were optimised for each gene, applying a minimum query coverage of 80% and identity thresholds as follows: 80% for psaA, 75% for hbsT (contigs: 65%), 70% for atpA (contigs: 60%), psbA (contigs: 60%), isoA, ygfK, and aro, 60% for amoA, mmoA, coxL, [FeFe]-hydrogenase, nxrA, rbcL, and nuoF, and 50% for all remaining genes (contigs: rho 30%; rdhA: 45%; cyc2: 35%). To estimate the relative abundance of community members harbouring each gene, read counts were normalised to reads per kilobase million (RPKM) as previously described45,46. Binned contigs were subjected to open reading frame (ORF) prediction using Prodigal v2.6.347. Resulting ORFs were annotated via DIAMOND blastp48 using the same reference database and identity thresholds as above, as well as in DRAM v1.5.049.

### Eukaryotic bins processing and metabolic annotation

Bins obtained from Vamb v4.1.330, MetaBAT v2.12.131, CONCOCT v1.1.032, and SemiBin v2.2.033 were filtered to a minimum length of 2.5 Mb, then contigs origin within each bin was estimated using Whokaryote v1.1.250 and only bins containing 80% or more eukaryotic contigs were retained. Bin quality was assessed with Busco v5.8.351 and those with 30% or higher completeness were retained. EUKulele v2.0.952 was used to infer contigs taxonomy within each bin. Bins were taxonomically classified when ≥70% of constituent contigs shared the same assignment. FastANI v1.3453 was used to dereplicate the bins at species level with 98% ANI according to their taxonomy, which after this step were referred as eukaryotic metagenome-assembled genome (eMAG). After these steps we retained two eMAGs, and calculated their abundance across samples using CoverM v0.7.040 in genome mode with method count. AUGUSTUS v3.5.054 with species “chlamy2011” was used to predict ORFs on eukaryotic contigs and eggNOG-mapper v2.1.1355 was used to annotate them against eggNOG v6.0 database56.

### Small subunit rRNAs reconstruction, classification and analyses

PhyloFlash v3.41^57^ was used to reconstruct small subunit (SSU) rRNA genes from quality filtered metagenomic short-reads generated using the BBDuk utility within BBTools v36.92 (https://sourceforge.net/projects/bbmap/), followed by targeted assembly of SSU rRNAs using SPAdes^58^ with standard configuration within phyloFlash. Reconstructed full-length and partial SSU rRNA sequences were taxonomically classified against the SILVA 138.1 SSURef NR99 database formatted for phyloFlash59. Read mapping was performed to quantify the relative abundance of Archaea and Bacteria across samples, resulting in a 16S rRNA gene-based abundance table used for downstream community composition analyses.

Community richness (α-diversity) was quantified using the Shannon index, and differences among sediment samples were assessed with analysis of variance (ANOVA) followed by Tukey’s Honest Significant Difference test. Community similarity (β-diversity) across sediment and seawater samples was evaluated from centered log-ratio–transformed Euclidean distance matrices of the 16S rRNA gene abundance table, after removing sequences with ≤10 counts, and visualised by principal component analysis (PCA). To test for differences in community composition among sediment samples and between sediments and seawater samples, we first assessed homogeneity of group dispersions using the ‘betadisper’ function in vegan60 and then determined statistical significance of between-group differences with permutational multivariate analysis of variance (PERMANOVA) implemented in the ‘adonis2’ function in vegan60. Linear discriminant analysis effect size (LEfSe)61, implemented in microbiomeMarker62, was applied to identify differentially abundant 16S rRNA gene sequences among entrance, midsection, and inner cave sediments. LEfSe analysis was performed at the genus level on the 16S rRNA gene abundance table after log10 transformation, with genera exhibiting LDA scores >2 considered indicative of enrichment within cave environments.

The multisite metric zeta diversity (ζ) was used to assess incidence-based turnover in community composition (16S rRNA gene sequences) across the cave environments using the zetadiv R package^63^. Zeta decline was calculated with the function ‘zeta.decline.mc’, performing subsampling set to 1,000 on a Jaccard-normalized presence-absence 16S rRNA gene dataset for ζ orders ζ1 to ζ3. This approach captured community structuring across the cave environments, with increasing ζ values approaching zero. Power-law and exponential models were fitted to the ζ decline curves, using Akaike Information Criterion (AIC) scores to evaluate which model best explained the relationship between ζ diversity and order *i*.

### *Ex situ* biogeochemical measurements

Incubation assays were carried out to assess aerobic metabolic activity within the cave communities. Each experiment was performed in triplicate, using three independent cave samples per environment (*n* = 9). Control samples were sterilised via autoclaving at 121□°C for 30 minutes, repeated twice. Elemental dynamics observed in live treatments were absent in sterilised controls, indicating that the measured changes were biologically driven.

Trace gas incubations were conducted in 120 mL glass serum vials containing approximately 10 g of sediment in 50 mL of autoclaved and 0.22 μm-filtered seawater, amended with 10 ppm of H_2_ and CO in the vial headspace. Gas sampling commenced immediately following the addition of electron donors, with 2□mL headspace gas extracted at defined time points. Gas concentrations were quantified via gas chromatography equipped with a pulsed discharge helium ionization detector (TGA-6791-W-4U-2, Valco Instruments Co. Inc.), using certified calibration standards (0, 10, and 100□ppm in N_2_; BOC Australia).

Nitrification incubations were conducted in uncapped 250□mL Schott bottles containing approximately 10 g of sediment in 100 mL autoclaved and 0.22 μm-filtered seawater, and supplemented with 100□μM NH_4_Cl. At each timepoint, 10□mL aliquots were filtered through 0.45□μm pore-size membranes and stored at -20□°C for subsequent analysis. Filtered samples were analysed for combined nitrite and nitrate (NO□^-^) concentrations using a Lachat QuikChem 8000 Flow Injection Analyzer (FIA), following standard APHA methods^64^.

For measurements of oxygen consumption, incubations were prepared in sealed 120□mL glass serum vials containing approximately 10□g of cave sediment in 100□mL of autoclaved and 0.22□μm-filtered seawater, and maintained in the dark while shaking. Dissolved O_2_ concentrations were measured at four time points over a 91 hours 25 minutes period using a FireSting oxygen probe (PyroScience).

Cell-specific oxidation rates were calculated for the samples used in the trace gas and nitrification incubations, by dividing bulk trace gas oxidation rates by the estimated abundance of organisms capable of each metabolism. Cell numbers were estimated from 16S rRNA gene copy numbers quantified by qPCR using a plasmid standard containing the 16S rRNA gene V4 region, adjusted by the average copy number of relevant metabolic marker genes per organism as estimated from short-read metagenomic data across the same samples, following a previously described protocol^46^.

### Radioisotope incorporation analysis

Cave sediment samples (0.5 g) with 1 mL of autoclaved and 0.22 μm-filtered seawater and an approximate concentration of 0.1 µM of radiolabeled sodium bicarbonate solution (NaH^14^CO_3_, Perkin Elmer, 53.1 mCi nmol^-1^) were prepared in 7 mL scintillation vials with ambient air headspaces. Quadruplicates of each sample were prepared for five experimental conditions: light (40 μmol m^−2^ s^−1^), dark + H_2_ (100 ppm), dark + CO (100 ppm), dark + S^2-^(800 µM), and dark + NH_4_^+^ (1 mM). Experiments were incubated for 3 days at room temperature, shaking at a constant rate (50-100 rpm). Following incubation, concentrated HCl was added dropwise to each vial and allowed to react for 24□h with intermittent shaking to ensure complete acidification and release of unbound dissolved inorganic carbon (DIC) as ^14^CO_2_. Acid was added uniformly across all samples until effervescence ceased before 350 µL of 10 M NaOH was added to neutralise. Finally, 5.2□mL of scintillation cocktail (EcoLume™, MP Biomedicals) was added and ^14^C activity was quantified using an automated liquid scintillation counter (Tri-Carb 2810 TR, PerkinElmer).

### Stable carbon isotope measurements

Stable isotope analysis was conducted to investigate pathways contributing to organic matter formation in the cave environment. Approximately 1□g of sediment was dried at 60□°C for 24□h, homogenised using a clean mortar and pestle, and further dried overnight at 60□°C. Samples were weighed into 8□x□5□mm silver capsules and decarbonated by repeated treatment with 10% HCl, followed by drying at 60□°C until effervescence ceased. Isotopic analyses were performed using an ANCA GSL2 elemental analyser coupled to a Hydra 20-22 continuous-flow isotope ratio mass spectrometer (CF-IRMS; Sercon Ltd., UK). Analytical precision for δ^13^C measurements was ± 0.2‰. Internal standards (sucrose, gelatin, and bream) calibrated against international reference materials (USGS40, USGS41, IAEA C-6) were analysed alongside samples to ensure accuracy.

### Chlorophyll *a* measurements

Chlorophyll was extracted by adding 9□mL of acetone to 5□g of cave sediment (four samples from the cave entrance and four from the inner cave), followed by vortex mixing and sonication in a high-power ultrasonic water bath for 5 minutes. Samples were then stored at 4□°C overnight to facilitate pigment release. The following day, 1□mL of ultrapure water was added to each tube prior to centrifugation at 2,000□rpm for 10 minutes. From the supernatant, 3□mL of the clarified extract was transferred into a cuvette, and absorbance was recorded at 665 and 750□nm using an Eppendorf BioSpectrometer. Chlorophyll *a* concentrations were subsequently calculated using the following equation:

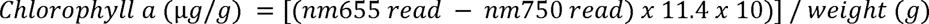

### Statistics and visualization

Downstream statistical analyses were performed in RStudio (version 1.2.5033) using R packages ggplot2^65^ for drawing plots and charts then Illustrator v24.0.2 was used for figure editing, phyloseq^66^, microbiomeMarker^62^, and vegan^60^ for community analysis, and ecolutils^67^ for Levins index^68^.

## Results & Discussion

### Physicochemistry and microbial structure change across a marine cave transect

Cave physicochemical conditions were profiled across a transect extending from the entrance to the innermost accessible environment. As we moved deeper into the cave, light levels dropped sharply towards total darkness **(Fig. 1c)**. In parallel, sediment grain size became finer (**Fig. S1**), suggesting lower hydrodynamic energy and transport from underwater currents. Sediment physicochemical analysis indicates a transition across the transect to a less alkaline (pH: 8.52 entrance; 8.41 inner) with lower organic carbon content (TOC: entrance 4.8%; inner 3.0%; **Fig. S2**) environment. The organic carbon within the cave likely reflects a combination of endogenous chemosynthetic primary production and lateral transport of photosynthetically-derived organic matter, balanced by consumption by organoheterotrophic microorganisms.

We tested whether archaeal and bacterial communities in the sediment varied across different cave environments and in relation to the overlying seawater. The innermost cave had the highest microbial richness despite exhibiting the lowest cell count (**Fig. 1d, e**). Microbial communities in the sediment differed significantly across the three cave environments and from the seawater (**Fig. 1f**). These observations (**Fig. 1d-f**), together with the clear distinction in physicochemical properties across the cave entrance, midsection, and inner cave (**Fig. S2**), suggest that deterministic processes may have been the leading force shaping the structure of the microbial communities colonising these environments. Consistently, *z-*diversity analyses^73^ showed microbial turnover across the cave environments and community assemblages were best described by a power law regression (AIC: -13.43; **Fig. S3**), indicating that they are likely structured by environmental gradients and exhibit habitat specialisation.

These inferences are reinforced by analysis of the relative abundance of microbial genera inhabiting the three cave environments (**Fig. 1g**; Supp. Data 1). Specifically, many taxa abundant in seawater were most abundant at the illuminated cave entrance, including diverse *Cyanobacteria* (**Fig. 1g**; Supp. Data 1), a result consistent with chlorophyll *a* quantification(**Fig. 1h**), and the highly abundant SAR11 clade (**Fig. 1g**; Supp. Data 1). Along the transect, the cave midsection showed the features of a transition zone with enrichment of taxa adapted to a broad range of redox conditions, from ammonia-oxidising bacteria such as *Nitrosococcaceae*^69^, through facultative anaerobes including *Reichenbachiella*^70^, to anammox bacteria “*Ca.* Scalindua”^71^ (**Fig. 1g**; Supp. Data 1). Moving deeper into the darker cave, enriched taxa comprise genome-streamlined putative fermenters such as *Woesearchaeales* and “*Ca.* Magasanikbacteria”^72^, as well as sulfide-cycling microorganisms such as *Desulfoconvexum*^73^ and *Sulfurimonas*^74^ (**Fig. 1g**; Supp. Data 1). Furthermore, the inner cave hosted taxa such as *Deferrisoma*, *Calditrichaceae*, *ABY1*, and *AT-s2-59*, typically associated with deep-sea habitats^75–78^ (**Fig. 1g**; Supp. Data 1).

### Functional traits distinguish cave generalists, entrance specialists, and interior specialists

To characterise the diversity, niche breadth, and functional potential of cave-dwelling microbes, we employed genome-resolved metagenomics, recovering 132 genomes spanning 14 phyla (**Fig. 2a**; Supp. Data 2). Most of the MAGs affiliated with bacteria, especially the *Proteobacteria* (59% MAGs; **Fig. 2a**), with one archaeal MAG (ammonia-oxidizing *Nitrosopumilus*; **Fig. 2a**) and two eMAGs (photosynthetic *Chlorophyta)* also retrieved (Supp. Data 2). We used Levins index^68^ to categorise each taxa into specialist, intermediate, or generalist based on their habitat distributions across the cave environments (**Fig. 2a**). Of the MAGs recovered from the sediment, 50% were classified as generalists found ubiquitously across the cave systems, whereas 28.8% of the specialist population colonised the lit cave entrance, and 10.4% the interior (Supp. Data 2). The habitat generalists may have flexible strategies to adapt to the varying physicochemical gradients within the cave ecosystem. Aligning with this concept, habitat generalists had larger genomes than specialists after normalising for genome completeness (4.6 ± 1.5 Mb *vs.* 3.7 ± 2.2 Mb; **Fig. 2b**) and substantially more genes (4,744 ± 1,466 *vs.* 3,858 ± 2,360; **Fig. 2b**), likely reflecting a broader repertoire of metabolic pathways and regulatory mechanisms. When examined separately, specialists from the cave entrance displayed reduced genome sizes (3.2 ± 2.2 Mb; **Fig. 2c**) and gene repertoires relative to those inhabiting the midsection and inner cave (5.1 ± 1.7 Mb; **Fig. 2c**). This pattern is consistent with genome streamlining, a strategy associated with oligotrophic, light-exposed marine environments^79^, such as the cave entrance, where selective pressures favour genomes that retain only the core metabolic functions.

**Figure 2.**
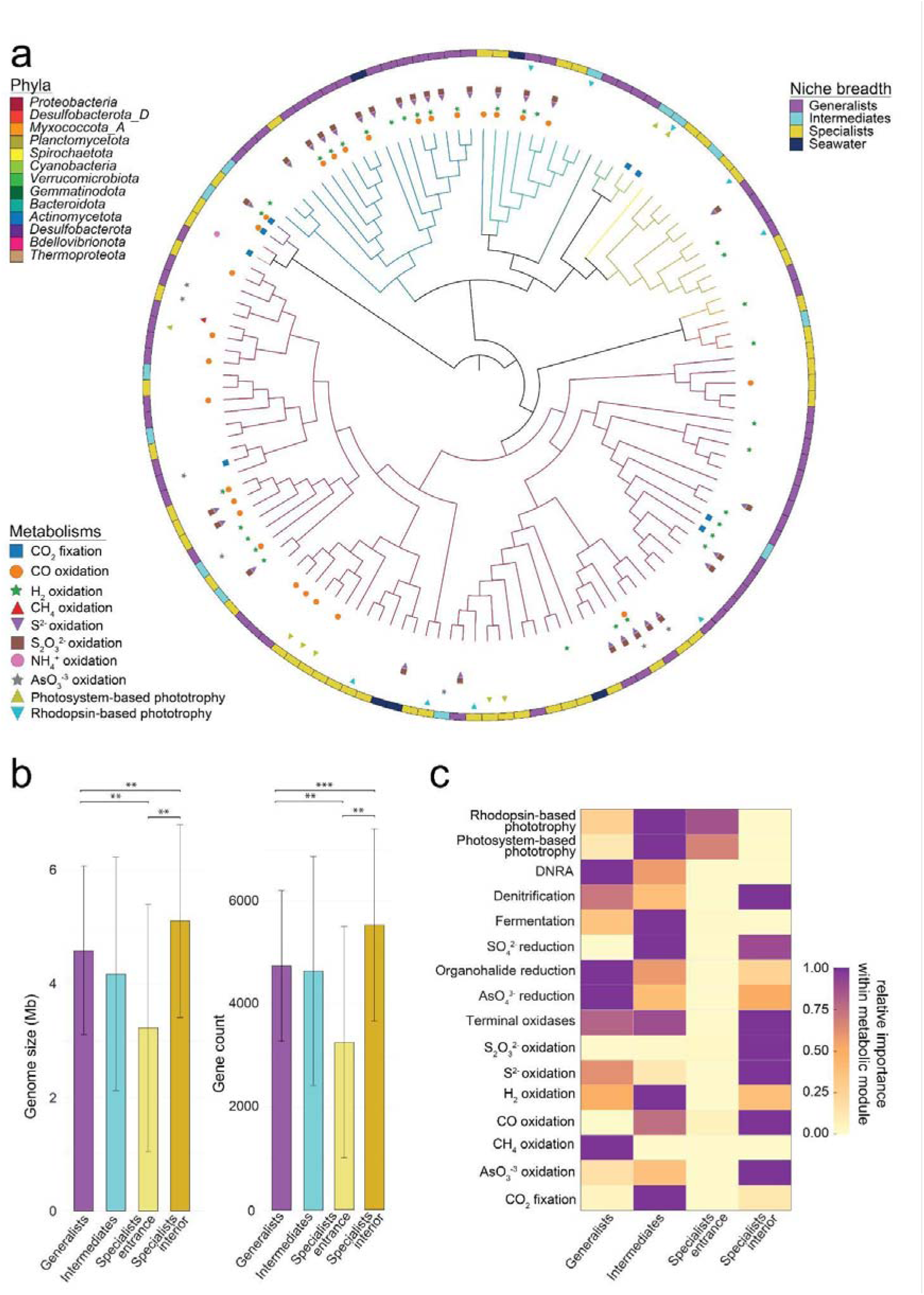
(**a**) MAGs maximum-likelihood phylogenetic tree inferred using LG substitution matrix, representing the topography, metabolic repertoires, and niche breadth of 130 archaeal and bacterial MAGs encompassing 13 phyla. Clades are coloured according to phylum-level taxonomy and symbols indicate the potential of each MAG to mediate key energy and carbon acquisition processes. (**b**) Bar graphs showing genome size (Mb) and gene count of generalists, intermediates, and specialists MAGs. Analysis of variance (ANOVA) shows statistical differences among generalists, intermediates, and specialists MAGs genome size and gene count (**c**) Metabolic functions of cave sediment MAGs and eMAGs identified as generalists, intermediates, and specialists.

The cave habitat generalists were predicted to be metabolically flexible microbes capable of using a wide range of energy sources, carbon sources, and electron acceptors (Supp. Data 2). Organoheterotrophy was pervasive, as highlighted by the diverse repertoire of carbohydrate-active enzymes (CAZymes), especially those involved in the breakdown of complex polysaccharides such as cellulose, chitin, and pectin, originating from plant, algal, and fungal sources (Supp. Data 2). This indicates that laterally transported organic carbon plays an important role in sustaining habitat generalists within the cave. In addition to using polysaccharides, some generalists were predicted to use one carbon compounds, as highlighted by the recovery of a putative methanotrophic *Rhizobiaceae* genome (**Fig 2a**; Supp. Data 2). This *Rhizobiaceae* genome encodes a XmoA-type copper-containing monooxygenase, but forms novel cluster distinct from characterised methane, ammonia, and hydrocarbon monooxygenases and thus its substrate is unknown (**Fig. S4**). Autotrophy emerged as a subsidiary trait among generalists and present in only 4.7% of this group, encompassing both photosynthetic and chemosynthetic microbes. The habitat generalist alga *Pycnococcus* was abundant throughout the cave, likely owing to recurrent deposition from seawater inflow where it is also common (Supp. Data 2), as well as its capacity to persist under low-light conditions, potentially supported by mixotrophic strategies such as heterotrophic carbohydrate scavenging (Supp. Data 2). Equally abundant were two chemoautotrophs classified as *Methyloligellaceae* and *Woeseiaceae* carrying genes for CO_2_ fixation via the Calvin–Benson–Bassham (CBB) cycle, supported by electrons harvested from the oxidation of S^2-^/ S_2_O_3_^2-^, and H_2_ respectively (Supp. Data 2). The capacity for lithoheterotrophy was also common, with more than 64% of generalist MAGs carrying genes for the oxidation of CO, H_2_, S^2-^, S_2_O ^2-^, and arsenite (AsO ^-3^; **Fig. 2a**; Supp. Data 2), with the capacity for H_2_ and S^2-^ oxidation 2.4- and 2.2-fold enriched in generalists relative to specialists (**Fig. 2d**; Supp. Data 2). Most generalists were predicted to be capable of both aerobic respiration (93% MAGs) and anaerobic respiration (80% MAGs), suggesting most are facultative anaerobes adapted to the spatiotemporal variations in oxygen availability in cave sediments. Particularly widespread was the capacity to use nitrate, nitrite, nitric oxide, nitrous oxide, arsenate, and organohalides as electron acceptors (Supp. Data 2). Fermentative metabolism was also ubiquitous, especially for H_2_ and acetate production (**Fig. 2d**; Supp. Data 2). Collectively, these findings indicate that cave habitat generalists can use a broad range of resources to thrive across the cave system despite sharp redox and resource gradients.

The habitat specialists inhabiting the lit cave entrance differed in their metabolic capabilities compared with those adapted to the cave interior. For instance, specialist facultative phototrophs including members of the *Rhodobacteraceae*, *Porticoccaceae*, *Pirellulaceae*, *Halieaceae*, *RS24* (*Alphaproteobacteria*), and *TMED25* (*Alphaproteobacteria*), as well as the alga *Picochlorum*, were most abundant at the entrance (**Fig. 2d**; Supp. Data 2), likely selected by increased light availability. In contrast, specialists inhabiting the midsection and interior were considerably enriched in genes for lithotrophic metabolism, including the oxidation of CO, H_2_, H_2_S, and S_2_O_3_^2-^ (**Fig. 2d**; Supp. Data 2), processes that can also sustain autotrophic carbon fixation. Consistent with this, members of the *Desulfocapsaceae* were abundant at the inner cave, where they likely exploit CO, H_2_, H_2_S as electron donors to fix carbon via the reverse tricarboxylic acid (rTCA) cycle (Supp. Data 2). The cave interior likely provides favorable environments for lithotrophic activity, where reduced hydrodynamic disturbance limits sediment mixing, promotes redox stratification, and fosters metabolic interactions in which anaerobes supply byproducts that fuel lithotrophs. The latter is reflected by the higher prevalence of genes associated with anaerobic metabolisms in interior specialists compared to entrance specialists; for instance sulfate reduction was exclusively encoded by this group, whereas denitrification, arsenate reduction, and organohalide reduction pathways were considerably enriched relative to the cave entrance (**Fig 2d**; Supp. Data 2). Altogether, these results indicate that specific functional guilds exhibit niche specialisation around spatially restricted energy sources.

### The inner cave environment is enriched in chemosynthetic communities

To test our hypothesis that chemosynthesis underpins cave microbial communities, we used gene-centric metagenomics to resolve how microbial metabolism shifted across the transect. Consistent with the physicochemical profiles and 16S rRNA gene analyses, the relative abundance of genes mediating CO_2_ fixation, trace gas metabolism, phototrophy, fermentation, nitrogen and sulfur cycling significantly varied across the cave environments (**Fig. 3a)**. Genes for photosystem- and rhodopsin-based light harvesting were present throughout the cave but decreased substantially in the inner cave, with photosystem II genes (*psbA*) declining by 76-fold (**Fig. 3a**; Supp. Data 3). In contrast, the inner cave environment hosted higher abundance of genes for the oxidation of inorganic compounds (**Fig. 3a**). [NiFe]-hydrogenases were particularly enriched (entrance: 25.0 ± 12.1%, inner: 85.2 ± 16.6%; **Fig. 3a**), suggesting H_2_ oxidation is a key process. Global surface oceans are supersaturated with H_2_^16,80,81^, providing a potential source to support hydrogenotrophic metabolisms in the cave ecosystem. However, a portion of the H_2_ oxidised within the cave likely originates from *in situ* biotic production via fermentative pathways and nitrogen fixation, with the genes encoding fermentative [FeFe]-hydrogenases, group 3 [NiFe]-hydrogenases, and nitrogenases also enriched in the inner cave (**Fig. 3a**). CO oxidation also emerged as a key metabolism, with anaerobic CO dehydrogenases (*cooS*) increasing 20.8-fold in the inner cave, whereas aerobic CO dehydrogenases (*coxL*) remaining relatively stable (**Fig. 3a**). Consistent with the likely presence of redox gradients across the cave, the inner cave was enriched with genes mediating anaerobic respiration, including key denitrification steps (*narG, napA*), dissimilatory nitrate reduction to ammonium (*nrfA*), and dissimilatory sulfate reduction (*dsrA*). Byproducts of these pathways, such as NO ^-^, NH□□, and S^2-^ from hypoxic niches, likely fuel chemosynthetic activity. Accordingly, genes for NO_2_^-^ (*nxrA*), and S_2_O_3_^2-^ (*soxB*) oxidation were detected across the whole cave (**Fig. 3a**), whereas S^2-^ (*sqr*) and NH_4_^+^ (*amoA*) oxidation genes were enriched in the inner cave (2.6- and 2.0-fold; **Fig. 3a**).

**Figure 3.**
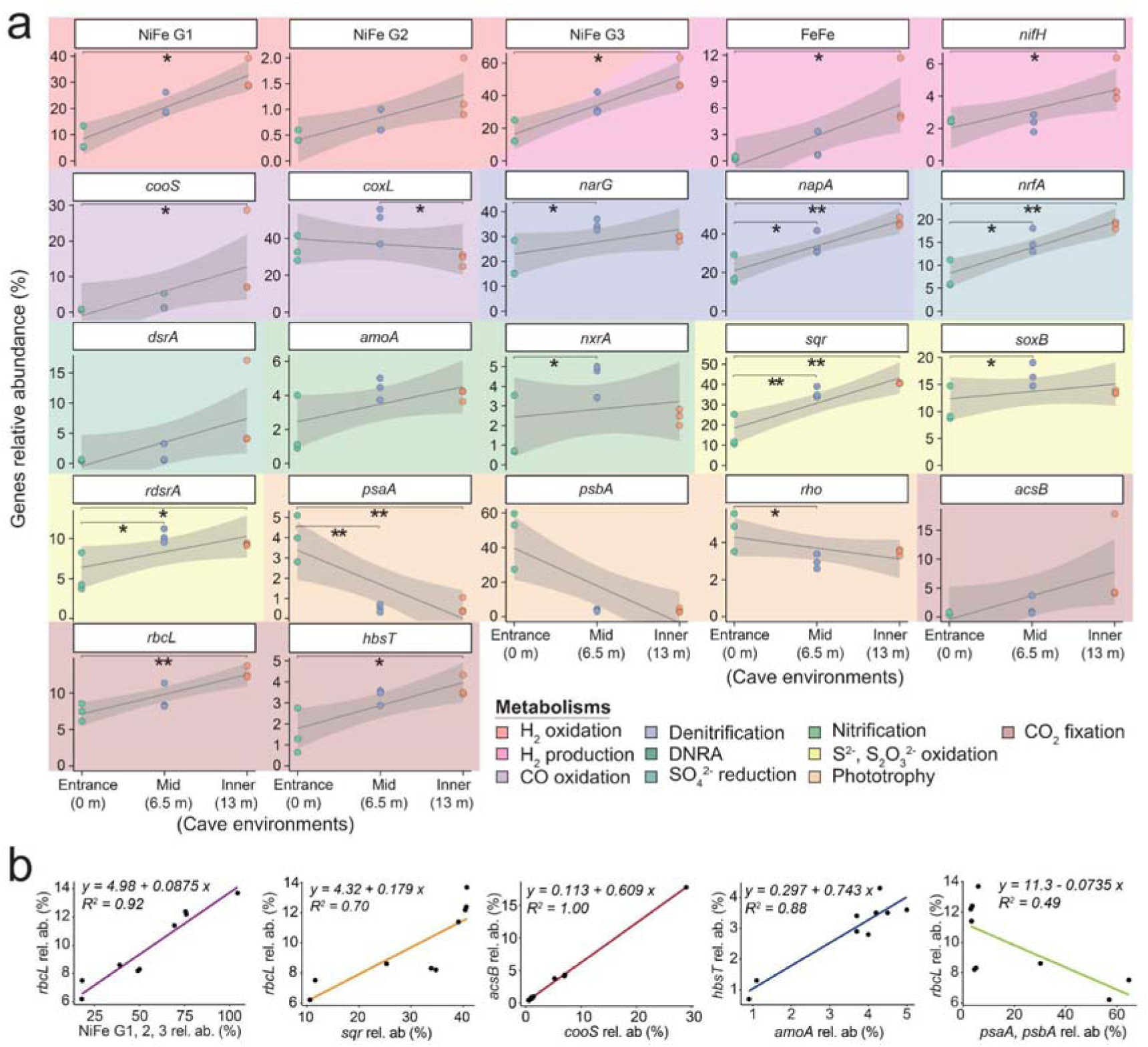
(**a**) Relative abundance of genes encoding enzymes for trace gas metabolism, fermentation, sulfur and nitrogen cycling, phototrophy, and carbon fixation across distinct cave environments. Significant differences in gene relative abundance across the cave environments were assessed using categorical models (ANOVA or Kruskal–Wallis test) with post hoc pairwise comparisons (Dunn’s tests). We also plotted regression lines across the transect distance (0, 6.5, and 13 m), which highlight the overall spatial trends. Shaded grey areas show 95% confidence interval of the regression lines. (**b**) Linear regressions showing correlations between the abundance of genes associated with energy metabolism (explanatory variable, x-axis) and those involved in carbon fixation pathways (response variable, y-axis) across the cave environments.

Potential for inorganic carbon incorporation through autotrophic pathways was high across the whole cave, but especially in the inner environment. Marker genes for all CO_2_ fixation pathways except the rTCA cycle were enriched in the inner cave (Supp. Data 3).The CBB (*rbcL* 9.8 ± 2.6% of the community), WLP (*acsB* 3.7 ± 5.5%), and 4HBP (*hbsT* 2.9 ± 1.2%) were the dominant CO_2_ fixation processes and exhibited on average a 1.7-, 16.1-, and 2.4-fold increase between the entrance and the inner cave (**Fig. 3a**; Supp. Data 3), respectively. Enrichment of carbon fixation pathways in the inner cave likely reflects that cave microbes are more limited by lateral organic carbon inputs, but also potentially have an elevated supply of inorganic energy sources, in their likely redox-stratified sediments. Linear correlation analyses demonstrated that CO_2_ fixation genes were most strongly associated with those for inorganic compound oxidation rather than light capture, i.e. chemosynthesis not photosynthesis (**Fig. 3b**). Specifically, H_2_ and to a lesser extent S^2-^ oxidation genes were most closely associated with CBB cycle genes (**Fig. 3b**), whereas CO and NH□□ oxidation genes were most associated with the WLP and 4HBP (**Fig. 3b**) respectively. These inferences are consistent with the MAG-based predictions (**Fig. 2a**) and wider culture-based studies on chemosynthetic microorganisms^82–84^.

### Chemosynthetic metabolisms are active across the cave environments

To assess the activities of cave microbes, we measured rates of inorganic substrate consumption, photosynthetic and chemosynthetic carbon fixation (**Fig. 4a-c**), and aerobic respiration (**Fig. 4d**) through *ex situ* assays using sediment collected from the cave entrance and inner environments. In line with the genomic predictions, microbial communities from both locations consumed oxygen, with the inner cave mediating 1.4-fold higher bulk aerobic respiration rates (entrance: 217 O_2_ nmol L^-1^ h^-1^; inner: 302 O_2_ nmol L^-1^ h^-1^; **Fig. 4d**; Supp. Data 4). Aerobic respiration is likely primarily driven by a combination of organic carbon derived from both *in situ* chemosynthetic primary production and laterally transported allochthonous substrates. The cave communities also consumed the inorganic energy sources H_2_, CO, and NH ^+^ aerobically, with comparable bulk consumption rates across the two environments (**Fig. 4a**). However, the normalization against cell abundance revealed the cave interior communities mediated significantly higher cellular rates of H_2_ and CO oxidation (2 and 1.8-fold higher than the entrance, respectively), whereas cellular nitrification rates were slightly lower than at the entrance (1.4-fold; **Fig. 4b**; Supp. Data 4). This pattern suggests that highly active chemosynthetic communities reside in the inner cave.

**Figure 4.**
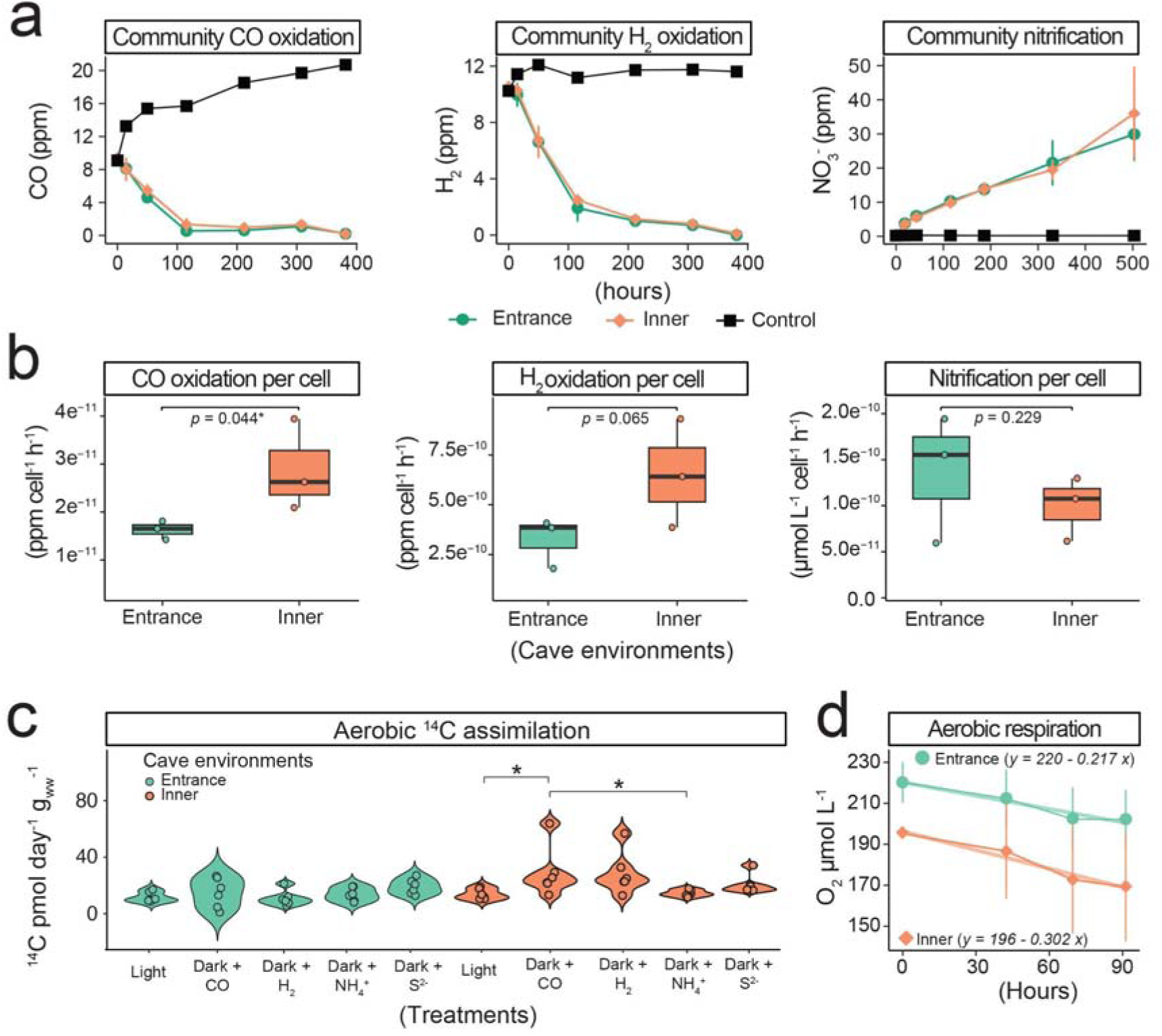
(**a**) Biogeochemical assays illustrating aerobic metabolic activity of microbial communities from the cave entrance and inner environments, based on the consumption of H_2_, CO, and NH_4_^+^. Heat-killed controls were included to confirm the biotic origin of the observed processes. (**b**) Oxidation rates of H_2_, CO, and NH_4_^+^ normalised per-cell specialised in each metabolism across entrance and inner cave environments. Shaded grey areas show 95% confidence interval of the regression lines. (**c**) ^14^CO_2_ incorporation by sediment community incubated under light (40 μmol m^-2^ s^-1^) and in the dark with CO (100 ppm), H_2_ (100 ppm), NH_4_^+^ (1 mM), and S^2-^ (0.8 mM). (**d**) Oxygen consumption rates of sediment from the entrance and inner cave environments. (**a**, **b**) Measurements were performed in 120 mL sealed serum vials containing ∼10 g of sediment and 50 mL (trace gasses) or 100 mL (O_2_ consumption) of 0.22 μm-filtered seawater. Trace gas incubations were supplemented with 10 ppm H_2_ and CO in the vial headspace. Nitrification assays (NO□ = NO_2_□ + NO_3_□) were conducted in uncapped 250 mL Schott bottles containing ∼10 g of sediment, 100 mL of 0.22 μm-filtered seawater, and 100 μM NH_4_□.

We complemented consumption assays with measurements of chemosynthetic and photosynthetic carbon fixation rates using radiolabeled carbon dioxide (^14^CO_2_). Incorporation of ^14^CO_2_ was consistently detected in all incubations, but as expected for sandy sediment collected at mesophotic depths^85^, photoautotrophy was minimal (**Fig. 4c**). At the cave entrance, chemosynthetic ^14^CO_2_ incorporation rates were low overall (14.1 ± 6.7 pmol^-1^ d^-1^ g_ww_^-1^) with modest stimulation by S^2-^ (19.2 ± 5.4 pmol^-1^ d^-1^ g_ww_^-1^) relative to other treatments (**Fig. 4c**; Supp. Data 3). In contrast, inorganic energy sources enhanced ^14^CO_2_ incorporation in the inner cave (**Fig. 4c**; Supp. Data 3). For instance, CO-supplemented incubations supported the highest ^14^CO_2_ incorporation with average rates 2-fold higher relative to the cave entrance (**Fig. 4c**). Likewise, H_2_-supplemented incubations incorporated on average 2.6-fold more ^14^CO_2_ in the inner cave compared to the entrance, consistent with the higher proportion of oxidative group 1 and 2 [NiFe]-hydrogenases (Supp. Data 3). The substantial stimulation by H_2_ in the inner cave aligns with observations from other subsurface and oligotrophic environments showing H_2_ is a key electron donor sustaining communities under nutrient- and light-limited conditions^86,87^. Conversely, NH_4_^+^ or S^2-^ did not stimulate ^14^CO_2_ incorporation, indicating that these compounds are either minimally used as electron donors by the communities or were already available in sufficient concentrations within the sampled sediment porewater (**Fig. 4c**). We note that our experiments likely underestimated the gross carbon fixation potential of the community due to the presence of abundant dissolved inorganic carbon in seawater and carbonates in the cave samples (**Fig. S1**), and the use of ^14^C-labelled CO_2_, which is discriminated against by microbial enzymes in favor of lighter, naturally occurring ^12^C.

The isotopic composition of bulk organic carbon (δ^13^C) was similar between the entrance (-22.5‰) and inner cave (-22.1‰) samples, though there is a slight enrichment in the inner cave (Supp. Data 4). The δ^13^C values nevertheless suggest organic matter in the cave was predominantly derived from CBB-based carbon fixation^88,89^. Since CBB cycle is the most abundant carbon fixation pathway in both cave photoautotrophs and chemolithoautotrophs, the exact contributions of photosynthesis and chemosynthesis to organic carbon supply in caves remain to be resolved, as does the relative contribution of allochthonous and autochthonous sources.

## Conclusions

Our findings demonstrate that even a relatively small marine cave harbours pronounced physicochemical gradients that structure microbial communities adapted to local conditions. Metabolically flexible generalists co-exist throughout the cave alongside specialists occupying well-defined niches shaped by resource availability; entrance communities rely on light-derived energy, either directly via phototrophy or indirectly through degradation of organic carbon (including phytodetritus), whereas interior specialists are increasingly dependent on inorganic compounds (either aerobically or anaerobically) for energy conservation and carbon fixation. The inner cave’s constant darkness, limited organic carbon inputs, reduced water flow, and redox gradients create unique pressures and a wide variety of niches for microorganisms. These conditions are ideal for chemosynthetic microbes that use inorganic compounds supplied to the cave or produced autochthonously by other microbes (e.g. sulfate reducers). Concomitant with decreased illumination, the progressively attenuated hydrodynamic flow along the cave transect likely promotes localised biogeochemical cycling that supports diverse metabolic processes. In particular, the reduced turbulence of the inner cave appears to facilitate the formation of stratified microbial communities, where chemosynthetic processes are fostered by compounds recycling between anaerobic and aerobic metabolisms. These patterns likely reflect ecological dynamics that have shaped the cave environments over time.

Our findings highlight that chemosynthetically-supported ecosystems are not confined to deep or extreme environments, but can form in aphotic habitats within the euphotic zone. Comparable gradients of light and hydrodynamics occur in many other marine settings, including caves, caverns, and crevices, but also artificial structures such as shipwrecks, solar farms, and piers, where chemosynthesis is also likely to play an important role in structuring ecological interactions and biogeochemical processes. Altogether, these results support emerging paradigms that chemosynthesis underpins a much larger fraction of oceanic productivity than was previously thought and reveals substantial hidden productivity in cryptic and littoral habitats.

## Data availability

All sequences generated from this work were deposited to the NCBI Sequence Read Archive. BioProject accession numbers for metagenomes and metagenome-assembled genomes are PRJNA1275038 and PRJNA1272733, respectively.

## Conflict of interest

The authors declare no conflict of interest.

## Acknowledgment

We are thankful to the boat skipper Neale Walker, the boat support team, and SCUBA divers Andrea Ceriani, Glyn Thomas, and Alex Hee Man Lee. We are also thankful to the Greening Lab members for inspiring and meaningful inputs. F.R. was supported by the Early Career Postdoctoral Fellowship (ECPF24-4273843556) awarded by the Faculty of Medicine, Nursing and Health Science at Monash University. P.M.L. was supported by ARC DECRA Fellowships (DE250101210). C.G. was supported by an ARC Future Fellowship (FT240100502). This study used the MASSIVE M3 supercomputing infrastructure.

## Author contribution

F.R. and C.G. conceived the study. Experimental planning and design were conducted by F.R., C.G., T.H., V.E., W.W.W., and P.L.M.C. Fieldwork was conducted by F.R., T.H. and J.Z. with the help of the boat crew and support divers. DNA extraction was performed by T.J. Gas chromatography measurements were carried out by F.R., T.H., J.Z., and T.N., whereas V.E. and W.W.W. oversaw nitrification measurements. ^14^CO_2_ assimilation incubations were performed by T.H. and J.Z. Stable isotope quantification was performed by W.W.W. Metagenome analysis, MAGs and eMAGs construction, and annotation were completed by F.R., with bioinformatics support provided by P.M.L. Formal analysis was performed by F.R. Resources, supervision, and funding were contributed by C.G. and P.L.M.C. The manuscript was written by F.R. and C.G., with input from all authors.

## Supplementary data caption

**Figure S1.**
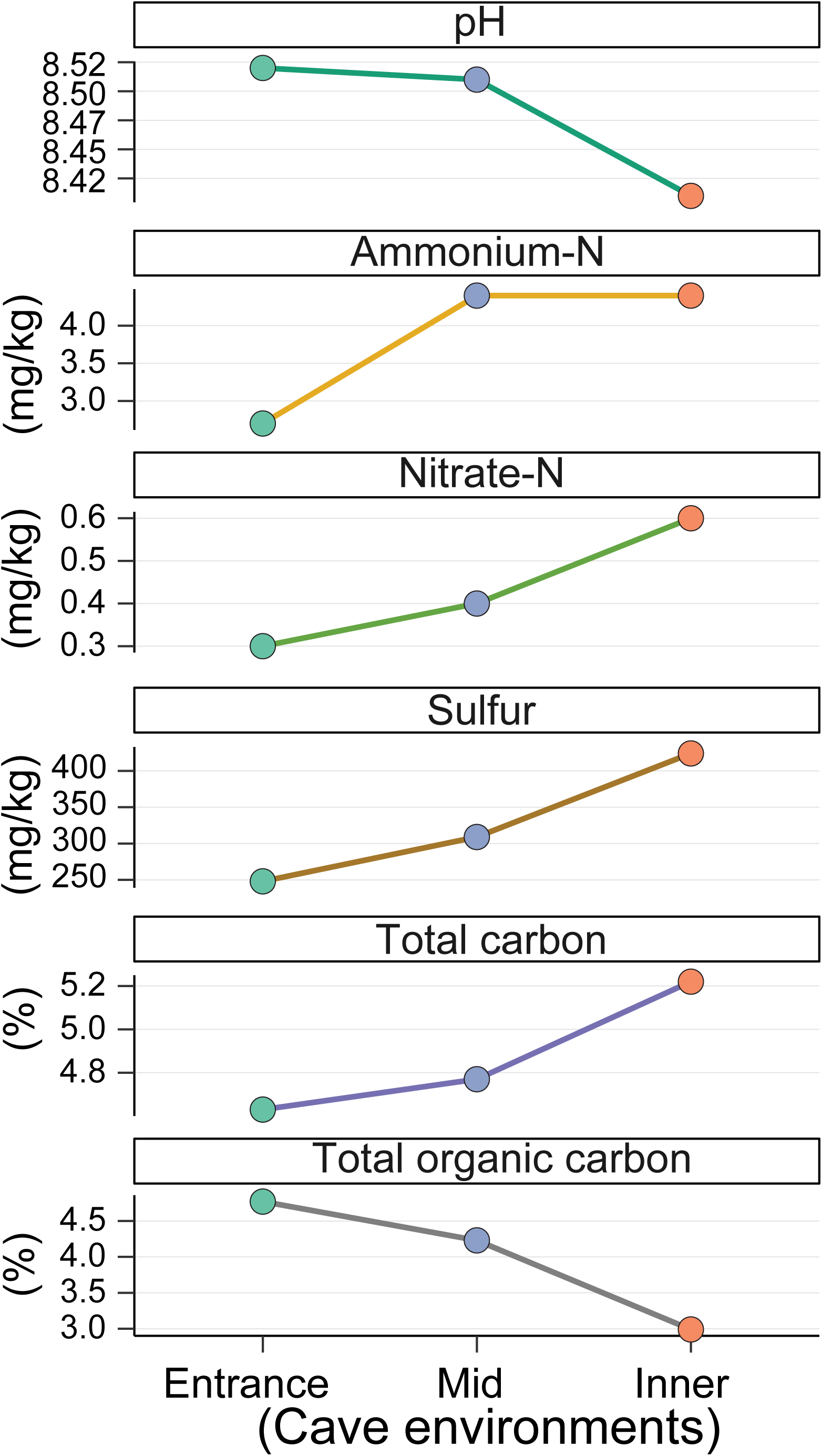
Sediment grain size from the cave entrance and inner environments.

**Figure S2.**
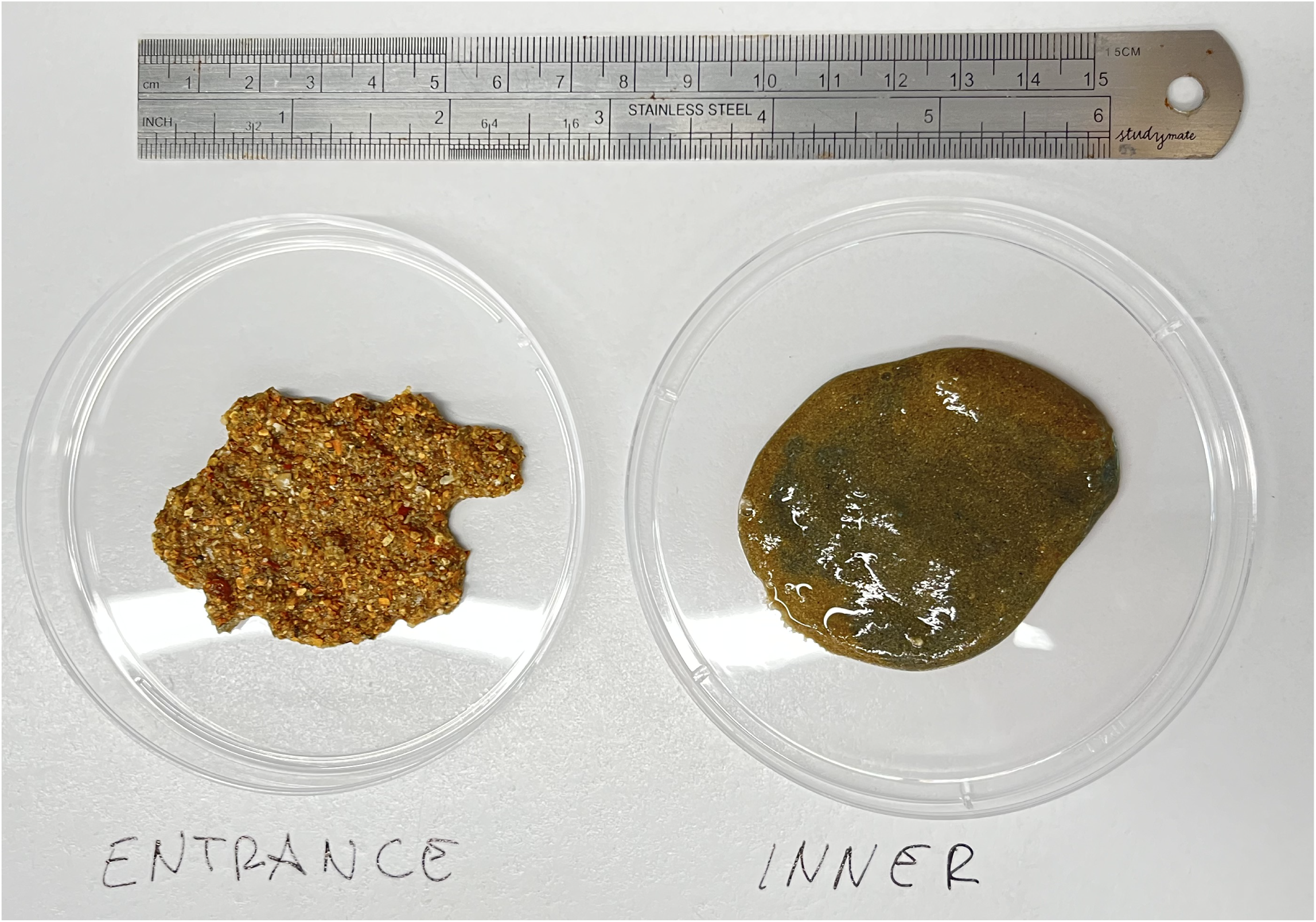
Sediment physicochemical properties across the three cave environments.

**Figure S3.**
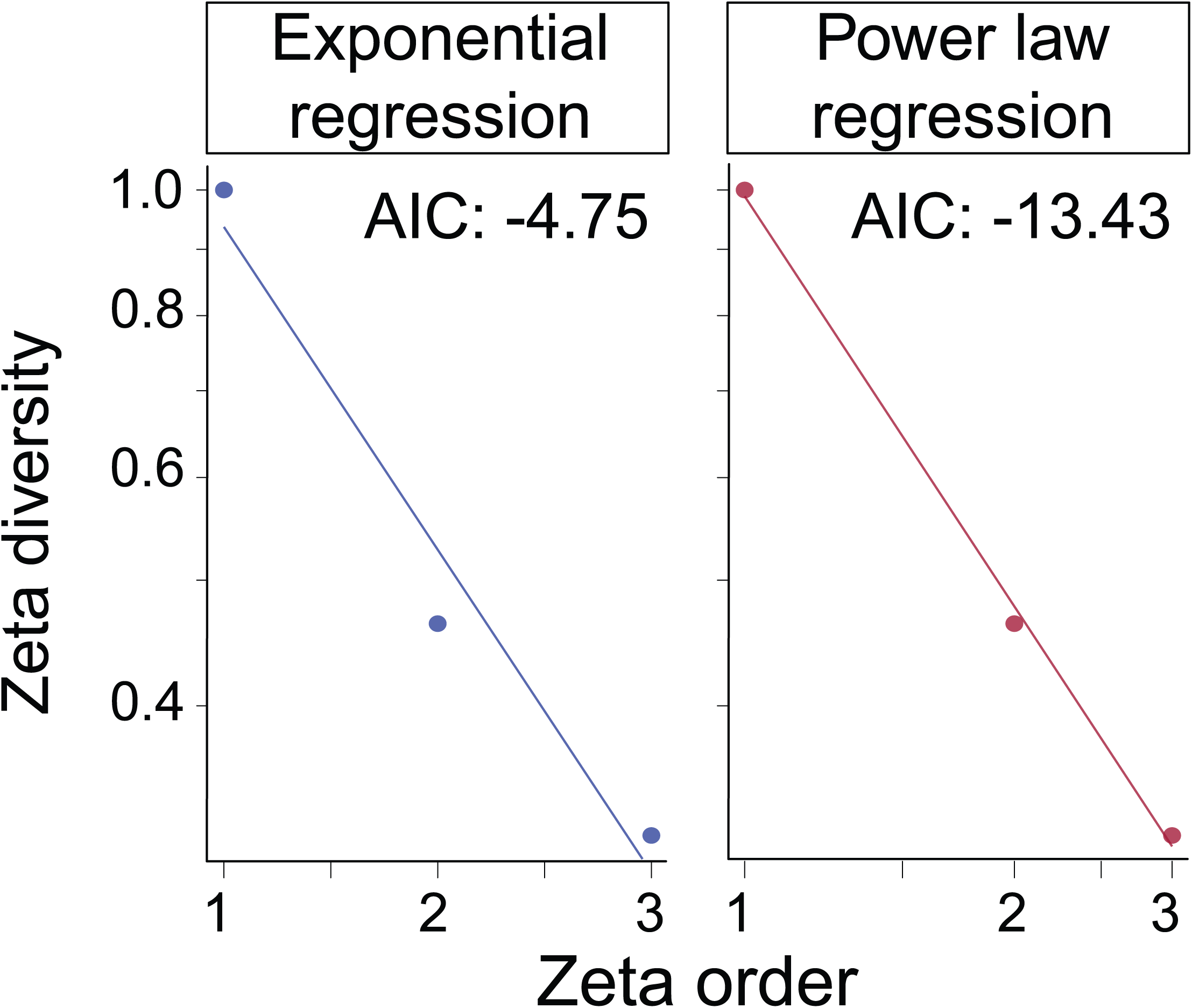
Zeta diversity (normalised Jaccard) decline along the sediment transect (entrance, midsection, innermost), modelled using exponential and power-law regressions, representing stochastic and deterministic processes, respectively. Model fit was assessed using Akaike Information Criterion (AIC).

**Figure S4.**
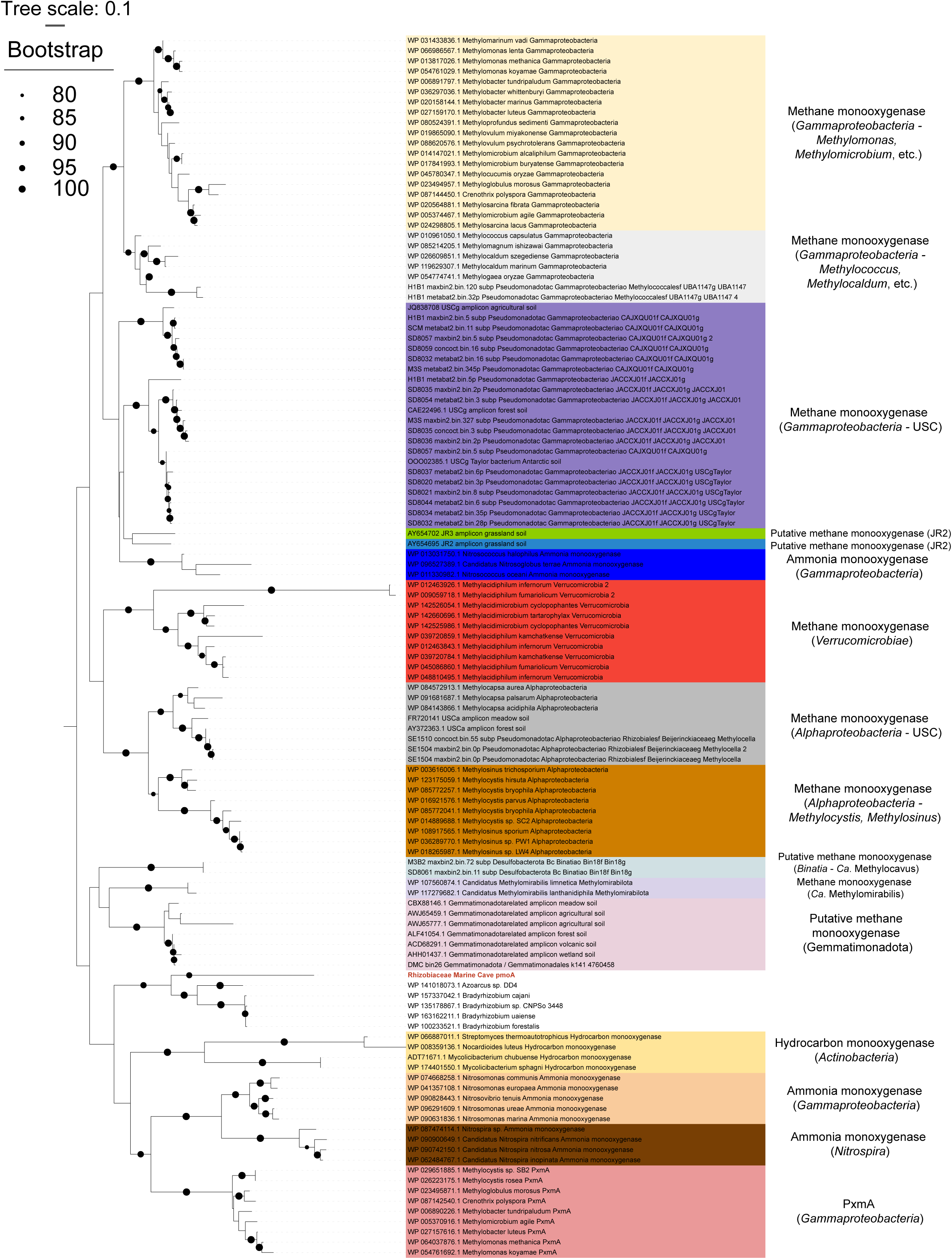
Maximum-likelihood tree of amino acid sequences of the alpha subunit of a copper-containing membrane monooxygenase (XmoA). The tree shows a sequence from a cave sediment metagenome-assembled genomes affiliated with the family *Rhizobiaceae* (bold red) alongside 118 representative sequences of the copper membrane monooxygenase superfamily, namely ammonia monooxygenase (AmoA), hydrocarbon monooxygenase (HmoA), and groups of unknown function (including PxmA). The tree was constructed using the LG+F+I+G4 model, bootstrapped with 1,000 replicates and midpoint-rooted.

**Supp. Data 1** (**a**) Sequence-based counts of bacterial and archaeal taxa across individual samples, based on taxonomic placement. (**b**) Cumulative counts of taxa across the cave entrance, mid, and inner environments. (**c**) LEfSe analysis identifying archaeal and bacterial genera enriched in entrance, midsection, and inner cave environments, with linear discriminant analysis (LDA) scores indicating discriminative power

**Supp. Data 2** (**a**) Features and abundances of metabolic marker genes associated with energy conservation, carbon fixation, phototrophy, and the cycling of hydrogen, carbon monoxide, methane, sulfur, nitrogen, and iron across 130 MAGs. (**b**) MAGs (n = 125) identified as exclusive to cave sediment. (**c**) Features and presence of carbon fixation and phototrophy marker genes in the 2 eMAGs. (**d**) Abundance of MAGs and eMAGs across the three cave environments. (**e**) Features and abundances of metabolic marker genes of MAGs with a chemosynthetic lifestyle. (**f**) Presence of carboxylases in sediment and seawater MAGs. (**g**) Presence of carbohydrate-active enzymes in sediment and seawater MAGs and eMAGs. (**h**) Presence of key fermentation enzymes in sediment MAGs. (**i**) Abundance of specialist, intermediate, and generalist MAGs across the cave environments. (**l-n**) Abundances of genes involved in carbon fixation, CO oxidation, H□ oxidation, CH_4_ oxidation, S^2-^ oxidation, S_2_O ^2-^, NH□□, phototrophy, and AsO ^2-^ cycling across specialist (by cave environment), intermediate, and generalist taxa, normalised to genome number and average completeness.

**Supp. Data 3** (**a**) Relative abundances of metabolic marker genes associated with energy conservation, carbon fixation, phototrophy, and the cycling of hydrogen, carbon monoxide, methane, sulfur, nitrogen, and iron, based on short-read metagenomic data from cave environments and seawater.

**Supp. Data 4** (**a**) H_2_, (**b**) CO, and (**c**) NH ^+^ cell-specific consumption rates in the cave entrance and inner environments. (**d**) Radioisotope incorporation (NaH_14_CO_3_) under treatments including dark, dark + CO, light, dark + S^2-^, dark + NH_4_^+^, and dark + H_2_ across cave environments. (**e**) Bulk organic carbon isotopic composition (δ^13^C) in samples from the cave entrance and inner environments. (**f**) Oxygen consumption measured over approximately 90 hours in sediment samples from the cave entrance and inner environments.

## Notes

### Competing Interest Statement

The authors have declared no competing interest.

